# Ketogenic diet administration later in life improves memory and regulates the synaptic cortical proteome through the cAMP/PKA signaling pathway in aging mice

**DOI:** 10.1101/2023.09.07.556691

**Authors:** Diego Acuña-Catalán, Samah Shah, Cameron Wehrfritz, Mitsunori Nomura, Alejandro Acevedo, Cristina Olmos, Gabriel Quiroz, Hernán Huerta, Joanna Bons, Estibaliz Ampuero, Ursula Wyneken, Magdalena Sanhueza, Felipe Arancibia, Darwin Contreras, Julio César Cárdenas, Bernardo Morales, Birgit Schilling, John C. Newman, Christian González-Billault

## Abstract

Aging is a complex biological process that compromises brain function and neuronal network activity, leading to cognitive decline and synaptic dysregulation. In recent years, a cyclic Ketogenic Diet (KD) has emerged as a potential treatment to ameliorate cognitive decline by improving memory in aged mice after long-term administration. However, the molecular mechanisms that govern such changes remains unclear. Additionally, whether short-term cyclic KD administration later in life preserves memory has not been addressed in detail. Accordingly, here we investigated how a short-term cyclic KD starting at 20-23 months-old regulates brain function of aged mice. Behavioral testing and long-term potentiation (LTP) recordings revealed that a cyclic KD improves working memory and hippocampal LTP in 24-27 months-old mice after 16 weeks of treatment. Moreover, the diet also promotes higher dendritic arborization complexity and dendritic spine density in the prefrontal cortex. Furthermore, to elucidate the molecular mechanisms underlying the memory improvements elicited by a cyclic KD, cortical synaptosomes of aged mice fed with this diet for 1 year were analyzed by mass spectrometry. Bioinformatics analysis revealed that long-term cyclic KD administration predominantly modulates the presynaptic compartment by inducing changes in the cAMP/PKA signaling pathway, the synaptic vesicle cycle, and the actin/microtubule cytoskeleton. To test these findings *in vivo*, synaptic proteins from cortices of 24–27-month-old mice fed with control or cyclic KD for 16 weeks were analyzed by western blot. Interestingly, increased Brain Derived Neurotrophic Factor abundance, MAP2 phosphorylation and PKA activity were observed. Overall, we show that a cyclic KD regulates brain function and memory even when it is administered at late mid-life and significantly triggers several molecular features of long-term administration, including the PKA signaling pathway and cytoskeleton dynamics, thus promoting synaptic plasticity at advanced age.

## Introduction

Age-related cognitive decline is the result of a complex process with multifactorial components that have a profound impact on the neuronal networks. Studies carried out in the brain of aged humans and rodents have identified a number of molecular alterations in the hippocampus and cortex, known as hallmarks of brain aging, that include mitochondrial dysfunction, impaired molecular waste disposal and aberrant neuronal network activity, among others (Mattson and Arumugam 2018). Such changes lead to defective neuronal activity characterized by dendritic tree regression (Burke and Barnes 2006), reduced dendritic spine density (Bloss et al. 2011), neurochemical imbalance (Bories et al. 2013), altered gene expression (Wruck and Adjaye 2020), loss of protein homeostasis (Morimoto and Cuervo 2014), and impaired glucose metabolism (Mosconi et al. 2008).

In the last decade, diverse strategies based on dietary interventions have led to improvements in learning and memory of aging mice. Remarkably, caloric restriction (Lin et al. 2015), intermittent fasting (de Cabo and Mattson 2019), fasting mimicking diets (Brandhorst et al. 2015) and ketogenic diets (KD) (Roberts et al. 2017) promote a metabolic switch that leads to fatty acid β-oxidation and consumption of ketone bodies instead of glucose for use as an energy source for neurons. A KD, which is a high-fat, low-carbohydrate diet, initially used as a therapy to treat refractory epilepsy (Lefevre and Aronson 2000; Sharma and Jain 2014) has been shown to not only enhance memory in older adults with mild cognitive decline (Krikorian et al. 2012; Mohorko et al. 2019) and preserve brain network stability (Mujica-Parodi et al. 2020), but also improve healthspan, memory and cognition in aging mice (Newman et al. 2017; Roberts et al. 2017; Zhou et al. 2021). This evidence demonstrates that a KD has neuroprotective effects on the brain, suggesting it has the ability to regulate synaptic function during the aging process. Indeed, β-hydroxybutyrate (BHB), the main ketone body produced in a KD, has a relevant influence on hallmarks of aging as it reduces the production of reactive oxygen species (Shimazu et al. 2013), enhances cell proteostasis (Finn and Dice 2005), activates mitochondrial biogenesis (Hasan-Olive et al. 2019), and prevents cell senescence (Han et al. 2018) and inflammation (Youm et al. 2015). Moreover, *ex vivo* and *in vitro* studies indicate that ketone bodies counteract neuronal hyperexcitability (Kadowaki et al. 2017), stimulate Aβ_1-40_ clearance (Versele et al. 2020) and activate autophagy in cortical neurons (Gomora-García et al. 2023). Additionally, the administration of ketone esters as an alternative way to deliver ketone bodies to the brain, improves cognition in humans (Fortier et al. 2021). However, despite compelling evidence showing the beneficial effects of a KD on brain function, numerous questions remain unanswered. For example, little is known about the molecular mechanisms that KD regulates at synaptic level and their effects later in life, whereas its impact on the neuronal proteome landscape and dendritic organization remains to be elucidated.

Therefore, we hypothesized that KD reconfigures neuronal networks in aged mice by regulating key proteins involved in synaptic activity, promoting enhanced long-term potentiation and neurotransmission. Indeed, using behavioral, electrophysiological, and biochemical approaches we found changes in critical proteins associated to synaptic function that will contribute to a better understanding of cognitive improvements that KD governs in the aging brain.

## Results

### Late mid-life cyclic KD administration increases β-hydroxybutyrate blood concentration and reduces glycemia in aged mice

In the first set of experiments, we defined a late life timeframe for the administration of the KD following a previous experimental setup (Newman et al. 2017). Specifically, 20-23 months old (m.o) mice were enrolled for a control diet (CD) or a cyclic ketogenic diet (KD) regimen for 4 months, as summarized in Fig. 1A. Such a strategy has previously shown that long-term non-obesogenic cyclic KD administration starting at middle-age prevents age-related memory decline in aged mice (24-26 m.o) (Newman et al. 2017), although whether the administration of this diet for a shorter length of time in late mid-life improves memory has not yet been addressed. We found that administration of a cyclic KD for just 4 months significantly increased β-hydroxybutyrate (BHB) blood concentration (Fig. 1B) and reduced glycemia (Fig. 1C) compared to the control group. Although body weight was higher in the KD group, the baseline before starting both diet treatments was greater for the KD mice compared to the CD group, a difference that was maintained throughout our study (Fig.1D, E). To minimize the stress of animal handling, all parameters were measured within the initial 12 weeks of treatment, before beginning behavioral testing.

**Figure 1.**
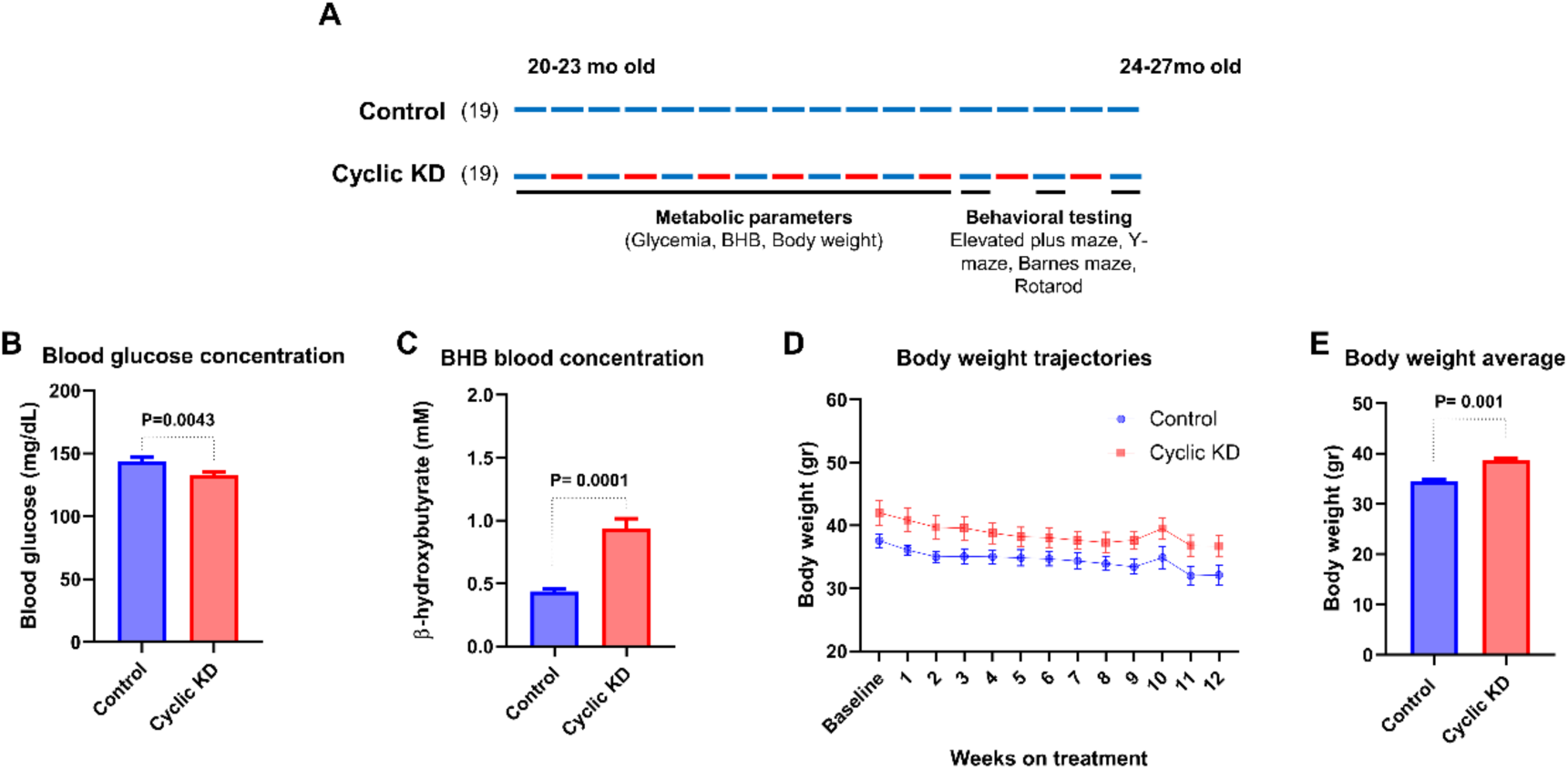
Control and cyclic KD regimens administered to aged mice for 16 weeks. (A) Scheme summarizing control and cyclic ketogenic diet administration, including measurements of metabolic parameters and behavioral testing. Each bar represents one week. (B-C) Glucose and BHB concentration were measured once a week by taking morning blood draws during the first 12 weeks of treatment (N=9). (D) Body weight trajectories. Even numbers represent KD weeks for the cyclic KD group, as indicated in red. (E) Body weight was measured at the beginning of the 12^th^ week of each diet regimen (N=14). Unpaired two tailed t test (p<0.005). Error bars show the SEM.

### Cyclic KD preserves memory and motor abilities after late mid-life administration in aged mice

To address the consequences of a KD on brain function during aging, treated mice performed a set of cognitive and physical tasks including the elevated plus maze, novel object recognition, open field, y-maze, Barnes maze and rotarod. To rule out confounding effects due to acute ketosis owing to administration of the KD, all tests were conducted in control weeks as shown in Fig.1A. Firstly, we assessed anxiety behavior using an elevated plus maze (Rodgers and Dalvi 1997) and found no differences in open arm entries between both groups, indicating that the food change did not induce anxiety in our cohort sample (Figure S1 A, B). Then, we analyzed locomotor and exploratory activity using the open field test (Seibenhener and Wooten 2015) revealing that KD-fed mice exhibited increased total movement compared to the control group (Fig.2 A). Interestingly, spatial recognition and working memory were preserved in aged mice fed with a cyclic KD, evidenced by increased spontaneous alternation, a condition that is unrelated to increased total movement (Fig.2 B and C). The Barnes maze test consisted in 4 consecutive training days, followed by an initial test on day 5 and a subsequent test on day 12. Cyclic KD group displayed a tendency to reduced latency across the days of training, specifically at day 4 (Fig.2 D). Although no differences were found at the testing day 5 (Fig.2 E), spatial learning and long-term memory were maintained in KD-fed mice, as shown by reduced latency to find the escape hole at day 12 (Fig.2 F). Importantly, as the Barnes maze test spans over two consecutive weeks, the second test on day 12 was conducted during KD week.

**Figure 2.**
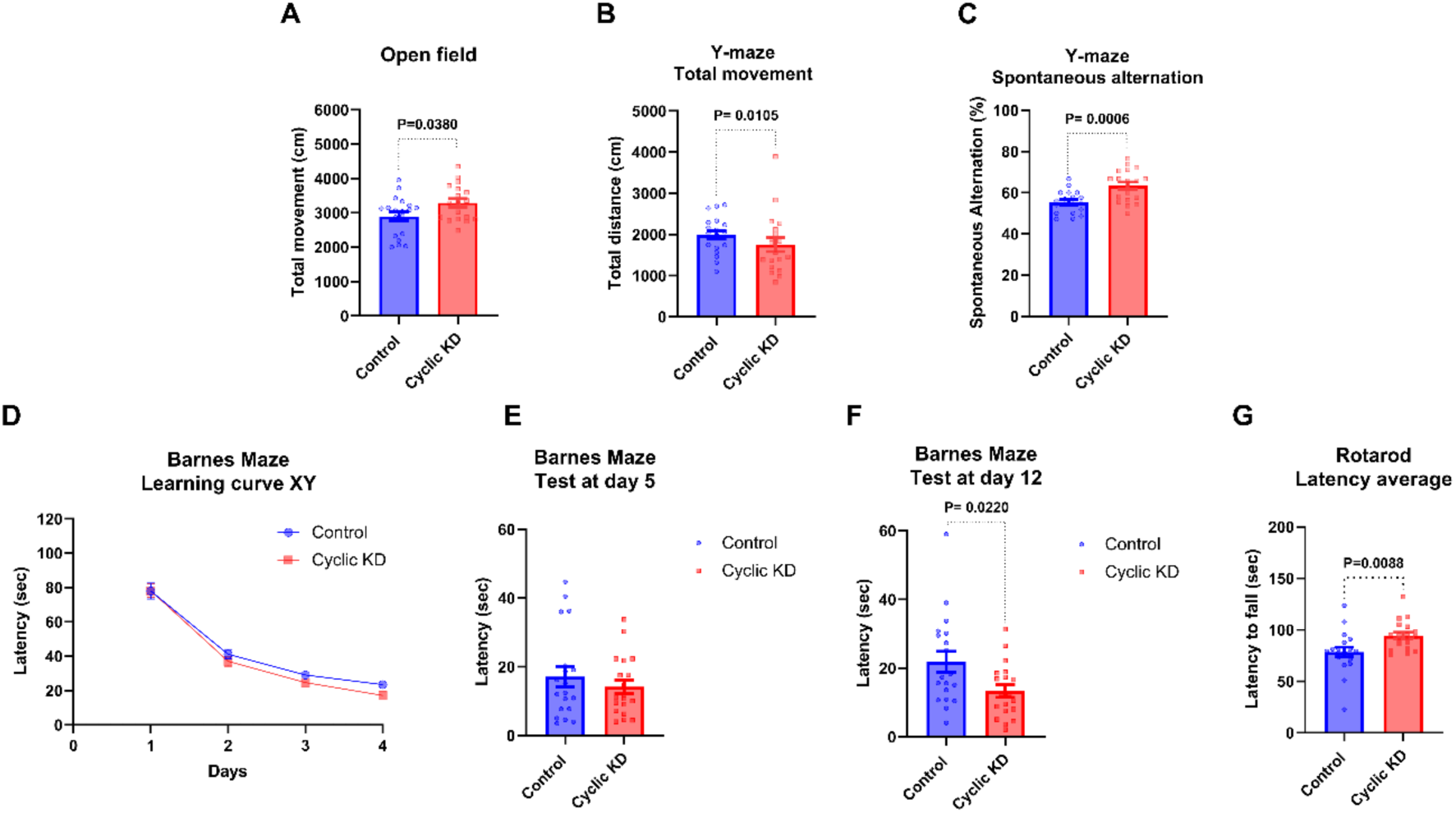
A cyclic KD administered at late mid-life improves memory and motor coordination in aged mice. (A) Total movement was calculated in the open field test. (B) Time that mice spent exploring new objects was indicated as percentage. (C-D) The Y-maze revealed increased covered distance and spontaneous arm alternation for the KD group. (E-G) The Cyclic KD group showed reduced latency to complete the Barnes maze test at day 12. (H) Increased latency to fall from the rotarod device was found for the cyclic KD group (N=19). Unpaired two tailed t-test (p<0.05). Error bars show the SEM.

Finally, we verified enhanced motor coordination in the rotarod test for the KD group. (Fig.2 G). Altogether, using an ample battery of tests encompassing behavior and locomotor functions, the findings strongly suggest that this nutritional intervention improves brain functions.

### Cyclic KD improves synaptic plasticity in the hippocampus of aged mice

The memory improvements evidenced in the cognitive tasks of aged mice fed with a cyclic KD suggest that this dietary intervention could influence neuronal activity in the hippocampus and cerebral cortex. Therefore, we analyzed synaptic connectivity in hippocampal slices of aged mice using electrophysiological recordings. Long-term potentiation (LTP) experiments at the CA3/CA1 synapse were performed, and field excitatory postsynaptic potentials (fEPSPs) were measured before and after induction in hippocampal slices from control- and cyclic KD-fed mice. fEPSP slopes show augmented TBS-induced hippocampal LTP in KD-fed mice brain slices (Fig.3 A). The potentiated response obtained with the KD diet was significantly higher than the age-matched CD-fed mice (compare red to blue traces), although milder than a young adult (P60) brain section (green trace). LTP quantification confirmed a 15% increase in LTP amplitude in the KD group compared to the control. Additionally, the LTP of brain slices from young mice (P60) was included as a control (Fig.3 B). Basal synaptic transmission was also measured by input and output recordings; no differences between groups were observed, ruling out that the increase in LTP were due to differences in postsynaptic depolarization caused by an altered basal synaptic strength. A trend to a reduced paired pulse facilitation was shown in the KD group (Fig. 3 C and D), consistent with an increase in neurotransmitter release probability. These quantitation experiments indicate that the LTP response elicited in the KD-fed animals partially reverse the reduced age-dependent plasticity in old animals, providing an electrophysiological basis that supports improved performance in the behavioral tests.

**Figure 3.**
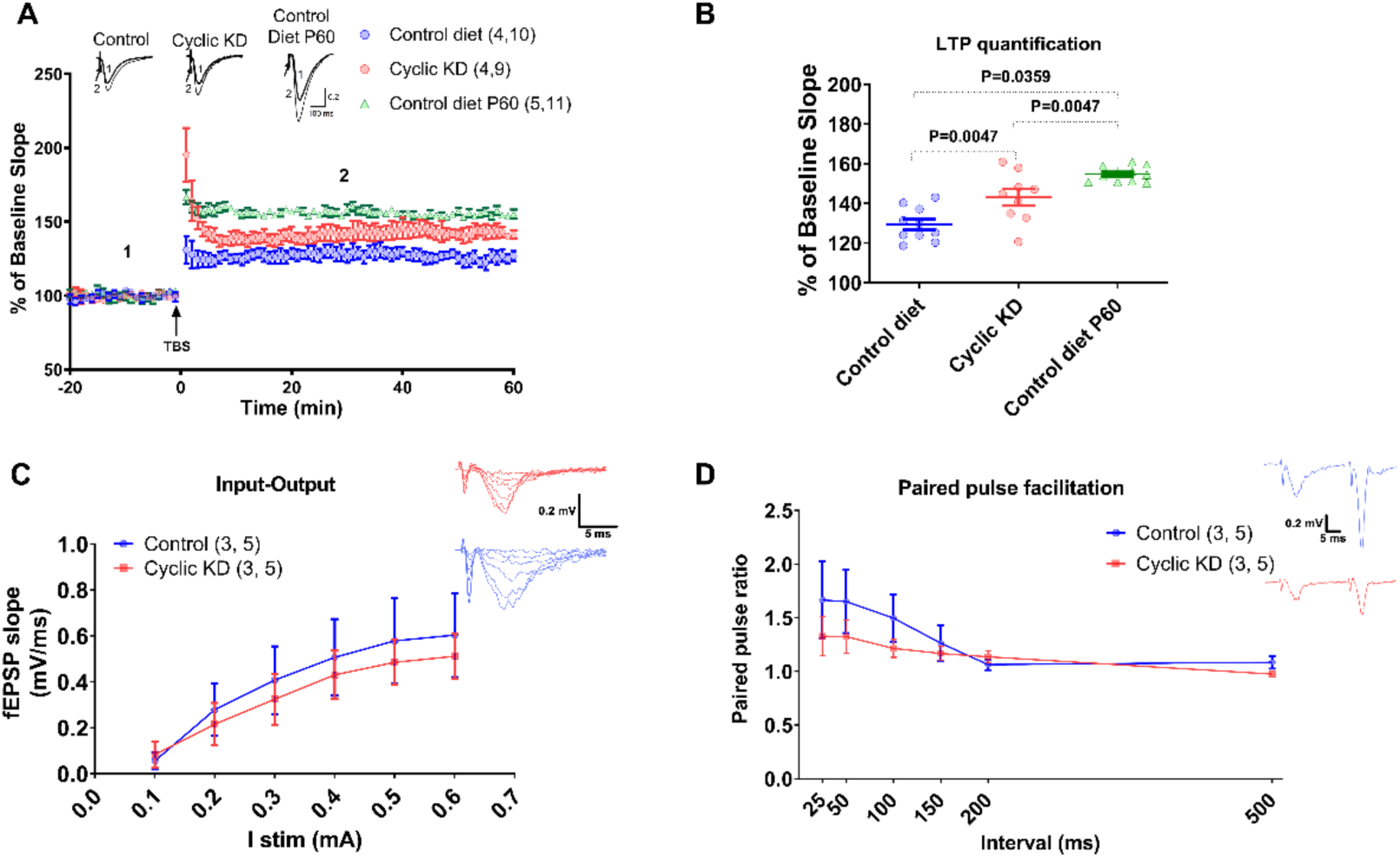
Hippocampal LTP is increased in aged mice fed with a cyclic KD after late mid-life administration. (A) fEPSP measurements expressed as normalized slope before and after TBS stimulation (black arrow) of slices from aged mice fed with control and a cyclic KD for 16 weeks. (B) Scatter plot illustrating the normalized fEPSP slope measurement assessed 50 minutes after TBS application. Hippocampal LTP response is significantly higher in cyclic KD-fed mice than in aged mice fed with a control diet (control: 129.3 ± 2.67, n=4.10; cyclic KD: 143.1 ± 4.19, n= 4.9; control diet (P60): 154.8 ± 1.15, n= 5.11; * p<0.05; ** p<0.01; **** p<0.0001; one-way ANOVA followed by Turkey post hoc test, F (2,27) = 22,26). (C) Basal synaptic transmission is similar in both cyclic KD- and control-fed mice. Average input-output curves obtained by increasing presynaptic stimulation are shown (2 animals per group; 5 slices for cyclic KD-fed and 4 slices from control-fed mice). (D) Paired pulse stimulation experiments at different interstimulus intervals. Summary plot of the fEPSP slope ratio revealing a trend for reduced paired-pulse facilitation in cyclic KD fed-mice (2 animals per group; 5 slices for cyclic KD-fed and 4 slices from control-fed mice). Input-output: Control: 0.8521 ± 0.2985, n=4; KD: 0.6892 ± 0.1065, n=5; p<0.05, t-student test. Paired pulse: multiple t-student test for each interval tested; p<0.05 in all cases.

### Cortical neurons from aged mice exhibit expanded dendritic tree complexity after administration of a cyclic KD at late mid-life

Electrophysiology assays revealed strengthened synaptic plasticity in brains of aged mice as a consequence of ingesting a cyclic KD. We reasoned that such changes should be accompanied by morpho-structural changes in neuronal morphology and/or the sites for synaptic contacts in dendrites. To determine whether changes in synaptic function was associated with rearrangements of synaptic organization, we assessed the morphological structure of neurons in aged mice. Golgi-stained brain slices from 26-27 m.o mice were imaged, including infralimbic and prelimbic areas of the prefrontal cortex -an area involved in working memory- to measure dendritic lengths of pyramidal neurons. A significant increase in arbor complexity was clearly evidenced in the cyclic KD group (Fig. 4 A). Sholl analysis confirmed such observations, as a robust increase in the number of intersections between 50-100 µm from the soma for the cyclic KD fed-mice was observed (Fig. 4 B). In the next set of experiments, we sought to analyze the number, distribution and shape of dendritic spines, the primary sites for excitatory neurotransmission in the brain. The morphology of dendritic spines is related with synaptic functional features. Therefore, we quantified the number of mushroom, stubby and thin spines in primary and secondary dendrites. We observed that in both CD and KD groups, stubby spines predominate over the other two classes. Like the structural changes observed in primary dendrites, we detected an increase in the total number of spines in animals fed with the KD (Fig. 4 C) and a significant rise in the number of stubby spines in primary dendrites derived from KD animals (Fig. 4 D).

**Figure 4.**
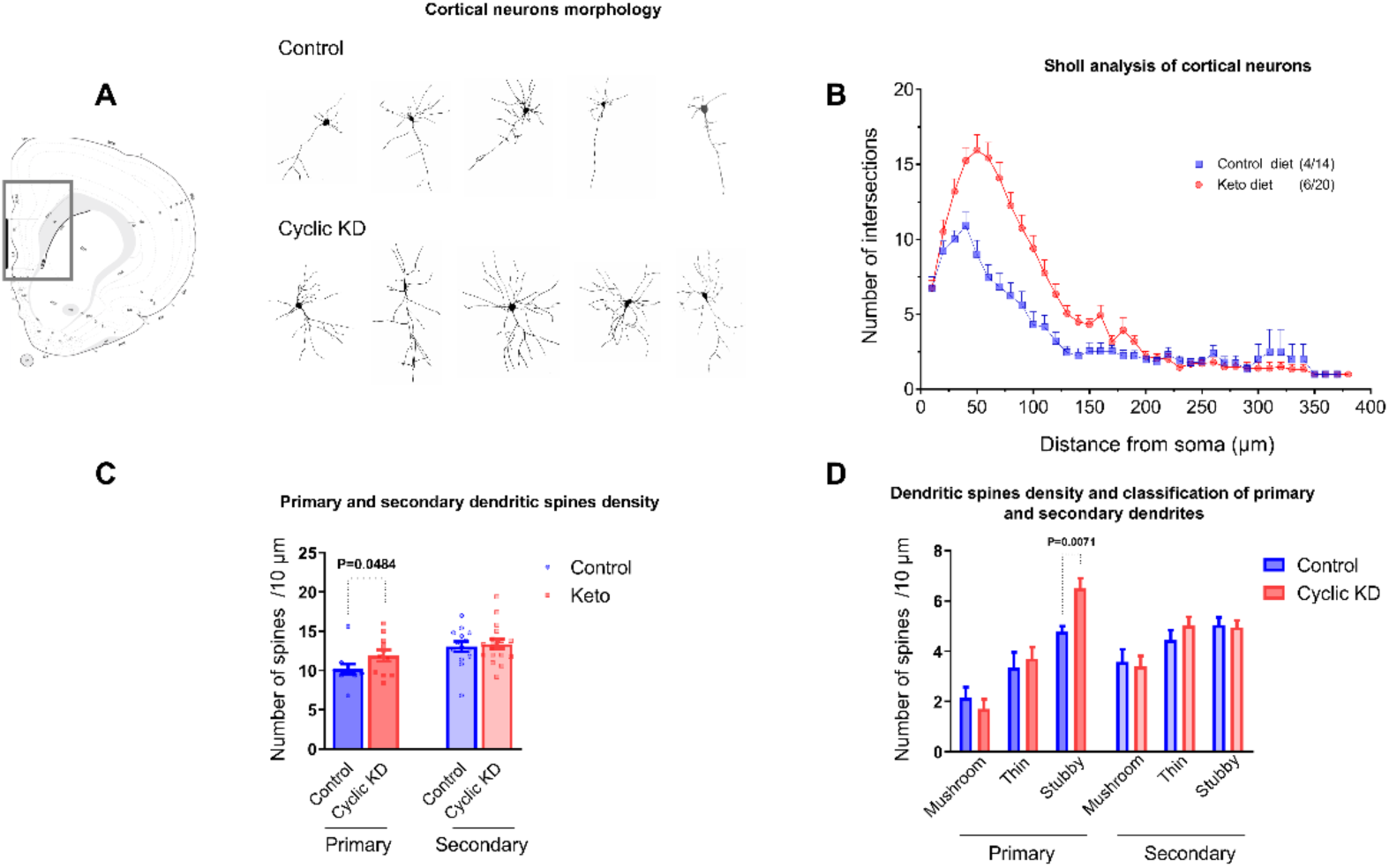
Cortical neurons from aged mice display higher dendritic tree complexity after a cyclic KD administration at late mid-life. (A) Cortical neurons (black rectangle) from brain slices of aged mice fed with control and cyclic KD were Golgi stained, drawn with camera lucida, and digitally scanned for morphological analysis. Representative images of pyramidal neurons from cortical layers 2/3 show dendritic tree arborization. (B) Sholl analysis comparing the number of intersections vs distance from soma between groups, revealed increased dendritic arborization in the cyclic KD group. (C, D) Dendritic spine density is enhanced in primary dendrites of the KD group whereas stubby spines are predominantly enriched in primary dendrites of KD-fed mice. Dendritic spine density: Control: 10.20 ± 0.6284; Cyclic KD: 11.90 ± 0.7460, unpaired t-test. Error bars show the SEM.

These experiments provide a morpho-structural basis for the significant recovery of neural plasticity properties in aged-animals observed in the electrophysiological assessments and suggest that the molecular machinery involved synaptic function might be differentially expressed in animals fed with the KD.

### Long-term cyclic KD administration regulates the cortical synaptic proteome through the cAMP-mediated signaling pathway, synaptic vesicle trafficking and actin cytoskeleton dynamics in aged mice

To gain a deeper understanding of how a cyclic KD influences synaptic function in the aging brain, we analyzed the cortical synaptic proteome of aged mice fed with a cyclic KD for 1 year (52 weeks; here referred as D52). The diet intervention started when mice were 12-14 m.o and it was administered in a cyclic manner for 1 year. Brain function was assessed by behavioral testing at 24-26 m.o according to (Newman et al. 2017). To avoid confounding effects of the ketosis state, brains were collected in a control week. Subcellular fractionation to obtain synaptosomes enriched in presynaptic and postsynaptic components from control- and cyclic KD-fed mice was performed according to (Dosemeci et al. 2006), including 4 biological replicates per condition. Successful separation and enrichment of synaptic components was evidenced by western blot, using SNAP-25, Synaptophysin, NR1 and PSD-95 antibodies as presynaptic and postsynaptic markers respectively (Fig S2. A, B) Subsequently, samples were analyzed by label-free data-independent acquisition (DIA) proteomics (Gillet et al. 2012; Collins et al. 2017; Schilling, Gibson, and Hunter 2017) which enabled the sensitive and accurate quantification of synaptic proteins by using tandem mass spectrometry (MS2) ion chromatograms. We quantitatively compared pre- and postsynaptic proteins derived from synaptosomes of aged mice fed with a cyclic KD versus CD. Significantly changed proteins (cyclic KD/CD) with ≥ 2 unique peptides, q-value<0.05 and absolute log_2_(fold change) ≥0.58 were considered (see Material and methods).

The unbiased proteomic characterization identified a total of 1782 synaptic proteins (see Supplementary InformationCandidates_DA3_old_0716_2020_v3all_proc1_0808_2023_SS.xlsx). Of those, 136 were differentially expressed with 123 presynaptic and 13 postsynaptic proteins. 69 proteins were up-regulated while 67 proteins were down-regulated when comparing cyclic KD versus CD (Fig. 5 A). To identify molecular targets regulated by a KD at the synaptic level, upregulated and down regulated proteins were separated to perform pathway and network analyses using ClueGO software (Bindea et al. 2009). Remarkably, we found a cluster of functionally related biological processes linked to actin dynamics, synaptic vesicle trafficking and cAMP-mediated signaling for the up-regulated proteins (Fig. 5 B and Fig. S3). On the other hand, mitochondrial function and electron transport chain were the most representative biological processes found in the down-regulated category (Fig. S4 A). Then, all differentially-expressed proteins of D52 were analyzed by SynGO (Koopmans et al. 2019), a specific database of functional synaptic gene ontology (GO). Notably, ‘synaptic vesicle’ was a GO term represented both in biological process and cellular component (Fig. S3 B, C). The synaptic proteins that were found to be present in SynGO biological processes and cellular component were included for protein-protein interaction analysis. The STRING database (von Mering et al. 2005) evidenced a strong association between actin dynamics related-proteins where Annexin V (Anxa5; postsynaptic), β-actin (Actb; postsynaptic), Wiskott-Aldrich syndrome protein family member 3 (WASF3; presynaptic) and Ras-related C3 botulinum toxin substrate 1 (Rac1; presynaptic) were the top up-regulated proteins in response to the KD treatment (Fig. 5 C).

**Figure 5.**
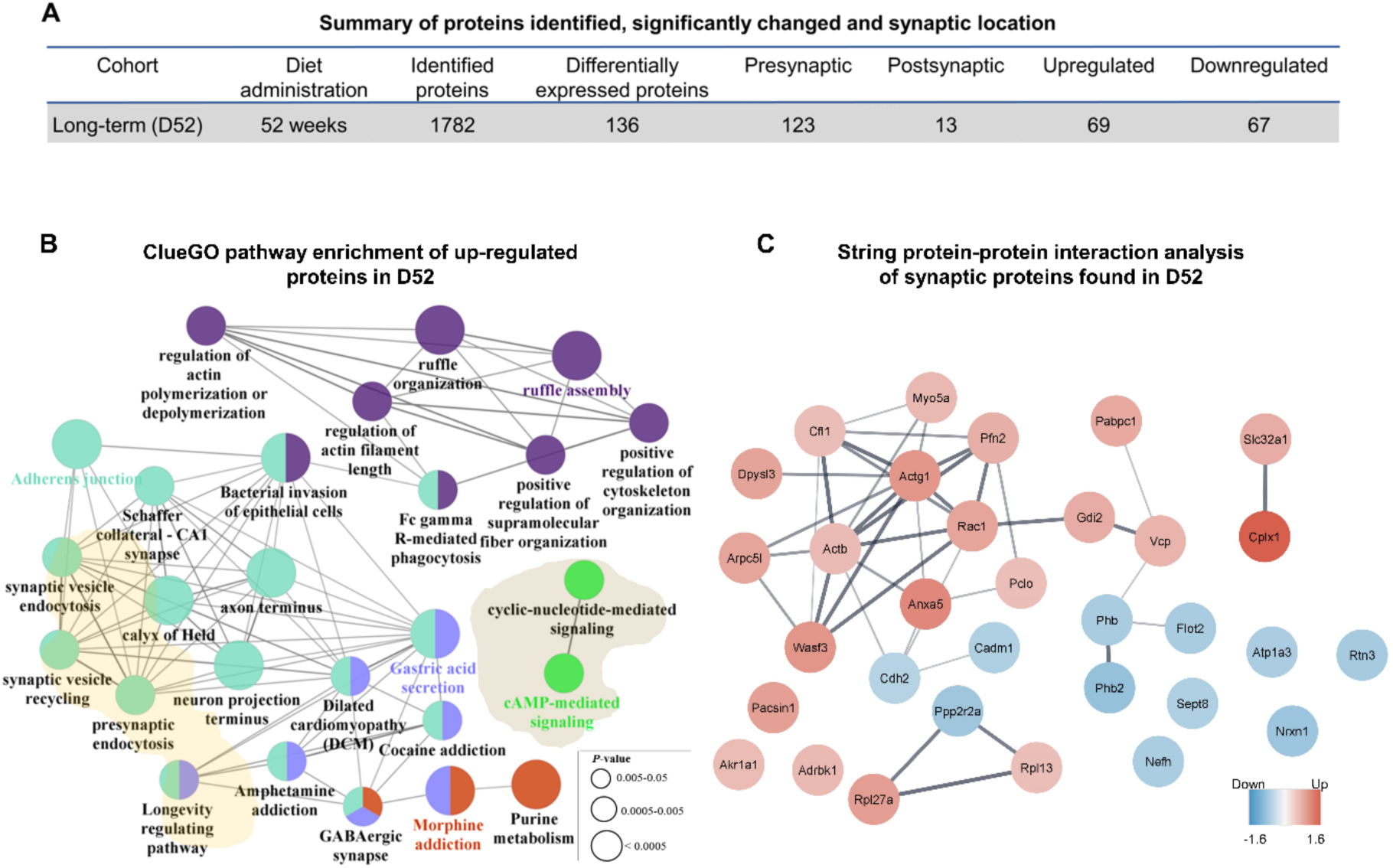
Long-term cyclic KD administration predominantly regulates the presynaptic cortical proteome through the actin cytoskeleton, synaptic vesicle trafficking and cAMP-mediated signaling in aged mice. (A) Summary of mass spectrometry analysis including total identified proteins, differentially expressed proteins and their subcellular location. (B) ClueGO pathway enrichment and network analyses of up-regulated proteins resulting from the cyclic KD/Control diet comparison. Pathways sharing the same color exhibit a similarity of at least 50%. Connecting lines indicate Kappa connectivity scores >40%. (C) STRING analysis based on protein-protein interactions from proteins contained within biological processes and molecular components derived from the SynGO database. Sphere color mapping is determined by the log_2_ ratio of the condition cyclic KD/Control diet. Thus, red indicates up-regulation and blue down-regulation. Each sphere is connected by a straight line, whose thickness represents the statistical power of the experimental evidence. The confidence score for interaction was 0.4.

Such an unbiased approach clearly revealed the up regulation of proteins that are related with improved neurotransmission such as those involved in actin dynamics and synaptic vesicles. In addition, these results suggest that the cAMP signaling pathway is specifically enhanced in animals fed with the KD.

### PKA kinase activity induces phosphorylation of downstream substrates in response to a cyclic KD and BHB administration in cortical neurons

Based on the bioinformatics findings, we decided to explore the cAMP-mediated signaling pathway in more detail. Cyclic AMP activates protein kinase A (PKA) which play a crucial role in neuronal function by regulating a number of protein phosphorylation events. Therefore, we scrutinized the activity of protein kinase A (PKA) by addressing the expression levels of phospho PKA substrates in prefrontal cortex of aged mice fed with a cyclic KD by western blotting. We took advantage of an antibody that recognizes specific p-PKA substrates phosphorylated in the RRXS*/T* motif (Muñoz-Llancao et al. 2017). Whole protein extracts of cortices from aged mice fed with a control or a cyclic KD were analyzed and probed with the p-PKA substrates antibody. A cyclic KD treatment promoted PKA kinase activity, as observed by the increased quantification of phosphorylated substrates with a electrophoretic mobility close to 130 KDa, 80 KDa and 55 KDa (Fig. 6 A and B). We also tested cAMP-pathway activation by-*in vitro* approaches. Interestingly an increased phosphorylation of a PKA substrate close to 130kDa was found in primary cultures of cortical neurons treated with β-hydroxybutyrate (BHB) for 1 hour, similar to observed *in vivo* (Figure 6 C). Additionally, CREB a transcription factor directly phosphorylated by PKA showed an increased p-CREB/CREB ratio in the soluble fraction after 1 hour of BHB treatment, which confirms the activation of cAMP-related signaling pathway in response to ketone bodies derived from a KD (Figure 6 C). Finally, to further support the idea that cAMP signaling increases with ketone body production, we evaluated *in vitro* changes in the activity of PKA, using AKAR4, a FRET biosensor (Depry et al. 2011). To this end, we transfected COS7 cells that were incubated with 5 mM BHB, and determined PKA activity. As shown in (Figure 6 D), 1 h of BHB treatment induced a robust increase in the YFP/CFP ratio, denoting increased kinase activity. Altogether these experiments show the increased expression of proteins relevant for brain functions and confirm the activation of the cAMP-signaling pathway in response to the KD treatment.

**Figure 6.**
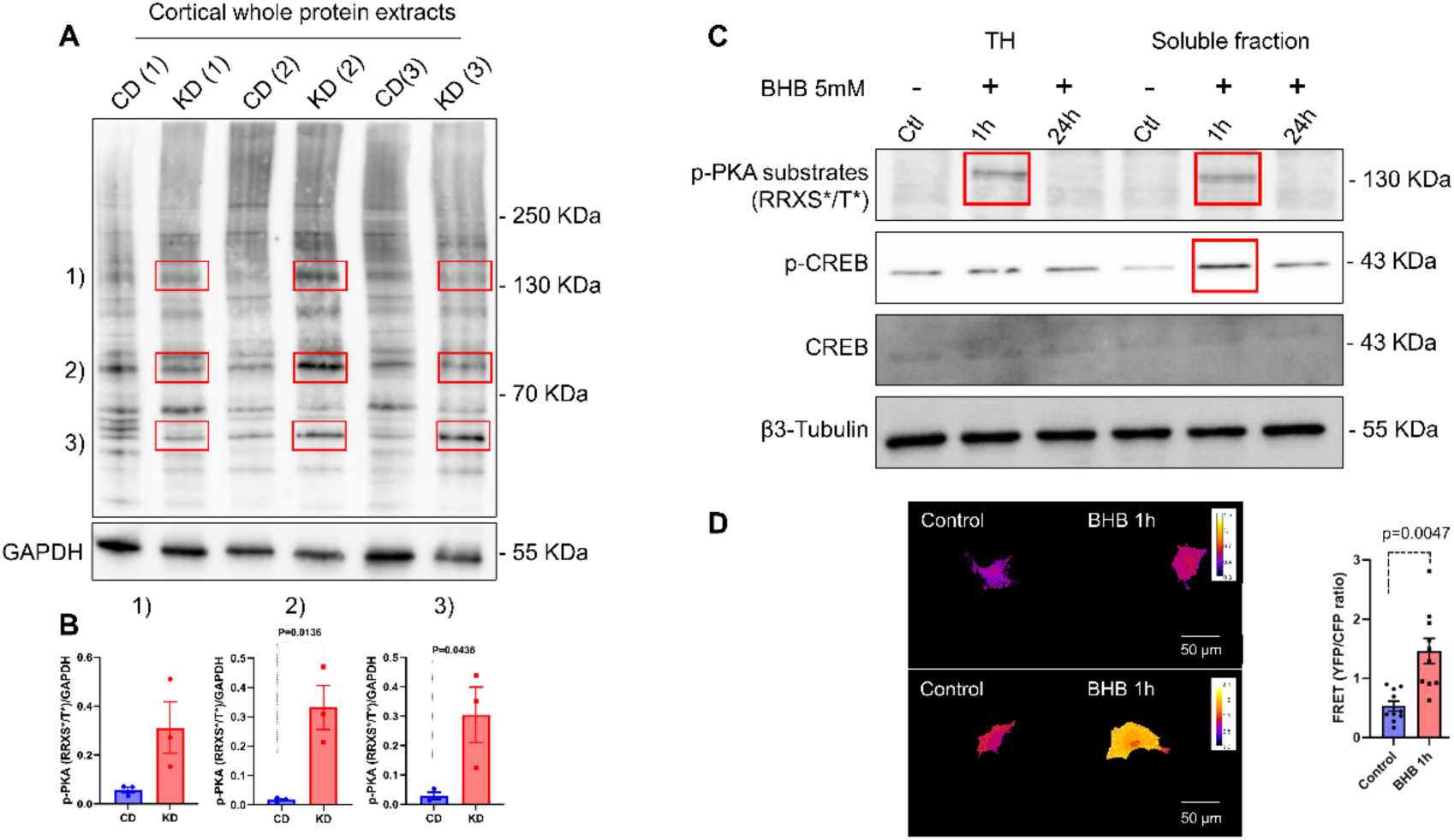
A cyclic KD promotes PKA kinase activity by substrate phosphorylation in the RRXS*/T* motif. (A) Whole protein extracts of the prefrontal cortex from aged mice fed with a control and a cyclic KD were analyzed by using a p-PKA substrates antibody that recognizes the specific RRXS*/T* motif. (B) Bands showing increased phosphorylation levels (highlighted in red boxes in A) were quantified (n=3 per group). (C) Total protein homogenates (TH) and soluble fractions from primary cultures of cortical neurons were treated with BHB for 1 h and 24 h and analyzed with p-PKA substrates antibody and pCREB/CREB antibody. (D) COS7 cells were transfected with AKAR4, a FRET biosensor for monitoring changes in PKA activity in response to BHB. Changes in protein expression are highlighted in red boxes. P<0.05, t-student test. Error bars show the SEM.

### A cyclic KD promotes BDNF expression and MAP2 phosphorylation in the prefrontal cortex of aged mice after late mid-life administration

Proteomics data collected from the D52 cohort revealed that cyclic KD activates the cAMP-related signaling pathway, which was confirmed by *in vivo* and *in vitro* studies of phospho-PKA substrates and CREB phosphorylation. Then, we decided to study the protein abundance of a set of synaptic proteins including BDNF, a canonical downstream target of PKA kinase activation. Consistently with the increased pCREB/CREB ratio western blots revealed elevated protein expression of BDNF (Fig. 7 A and B). Although the up regulation observed for PKA itself was not significant, an increased trend in the KD condition was noted (Fig. 7 A and C). Interestingly, microtubule associated protein 2 (MAP2), a protein associated to dendritic organization in neurons was found to be highly phosphorylated in the KD group (Fig. 7 A and D).

**Figure 7.**
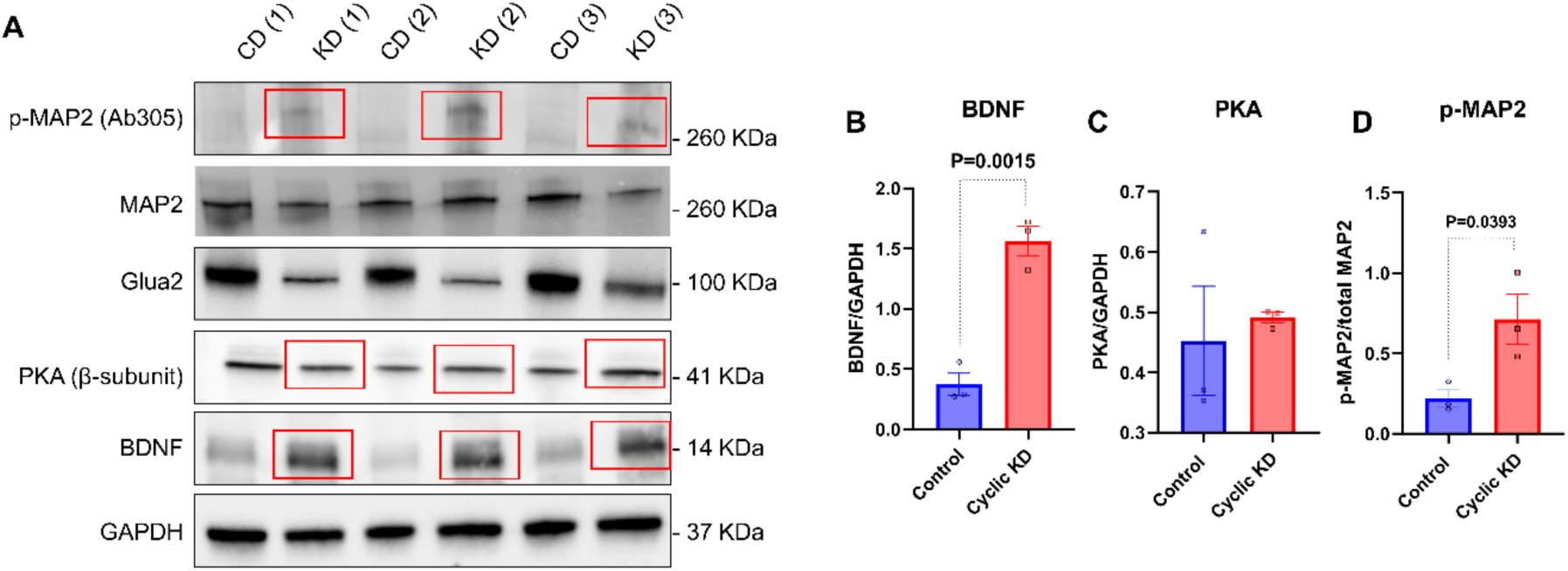
16 weeks of cyclic KD administration at late mid-life increases BDNF expression and p-MAP2 phosphorylation in the prefrontal cortex of aged mice. (A) Prefrontal cortices of aged mice fed with a control or a cyclic KD were collected in control week and processed to obtain whole protein extracts. Samples were analyzed by western blot (3 biological replicates) and probed against a set of synaptic-related antibodies. (B, C, D) BDNF expression was highly marked in the KD group, PKA catalytic subunit displayed a tendency to increase in response to KD, and p-MAP2 phosphorylation was increased in the KD condition (red boxes in A). Error bars show the SEM.

## Discussion

In this work, we provide cellular and molecular mechanistic evidence that an intermittent ketogenic diet (KD) in aged animals improve brain functions. Using behavioral, electrophysiological, morpho-structural, proteomic, biochemical, and cell-molecular approaches, we show that KD improves memory and potentiate synaptic function, remodels the synaptic proteome and activates PKA signaling. The activation of the cAMP-dependent signaling connect the different layers here studied, providing a novel mechanism for the beneficial effects observed after KD administration in aged mice.

Studies in the field of aging have consistently demonstrated that a KD reduces midlife mortality and modifies brain function in mice after long-term administration (Newman et al. 2017; Roberts et al. 2017). Such studies typically initiated diet changes starting at the middle stage of the animaĺs life, and evaluated the consequences later in life, after more than a year of diet administration. However, whether a KD can influence cognition and memory later in life, when the risk of age-related cognitive decline increases, remains to be elucidated. Recently, a study published by Zhou and cols. (Zhou et al. 2023) showed that ingesting a KD, starting at 18 months of age improves spatial memory and muscle endurance after 5 months of administration. Here, we report that 4 months of a cyclic KD administration (alternated weekly with a control diet to prevent obesity) starting at 20-23 m.o significantly improves working memory and long-term memory in 26-27 m.o mice (Fig. 2 D, G). Evidence of the effects of a KD on cognitive decline has been previously demonstrated. For instance, in an Alzheimeŕs disease model, a KD rescued spatial working memory of 5XFAD mice after 4 months of treatment, with no differences in locomotor activity (Xu et al. 2022). Interestingly, the same study showed that when the severity of cognitive decline is pronounced, the diet was not effective, as the administration of a KD to 5XFAD mice at a later stage (9 m.o) did not yield positive effects on brain function. Thus, studies concerning the effects of a KD on cognitive performance in normal aging at an advanced age are still limited and require dedicated attention. Our results demonstrate that in addition to memory improvements, a cyclic KD administered later in life also promotes increased motor coordination, as observed in rodent models with decreased locomotor activity (Shaafi et al. 2016; Streijger et al. 2013).

Synapses play a critical role in learning and memory processes. During aging, synaptic transmission efficiency declines both in the cortex and hippocampus with dramatical morpho-structural changes associated with decreased synaptic contacts, reduced dendritic complexity and altered dendritic spine dynamics (Morrison and Baxter 2012; Bloss et al. 2011; Jacobs, Driscoll, and Schall 1997; Griffin et al. 2006; Rosenzweig and Barnes 2003). Here we report that a cyclic KD restores LTP in the hippocampus of aged mice, as it was significantly improved compared to the control group. Moreover, we included LTP measurements of P60 mice as a young control, suggesting that cyclic KD reverses LTP impairment with age, resulting in a performance closer to young mice values. Interestingly, this was consistent with the reduced latency to find the escape hole in the Barnes maze, 12 days after the test was initiated, which supports a role of a KD in long-term memory. To our knowledge, this is the first time that this finding has been reported in mice of such an advanced age (26-27 m.o). Input-output curves obtained from fEPSPs recordings did not change between groups, which suggests no differences in basal transmission; nevertheless, the KD group showed a tendency to reduce paired pulse facilitation which could be associated with an increased probability of neurotransmitter release, which might contribute to a more efficient LTP induction by TBS.

Our morphometric analysis revealed higher dendritic arborization in the KD group in the 50 µm closest to the cell body, a feature that was correlated with an increased dendritic spine density in primary dendrites. In contrast, reduced dendritic complexity has been found in the medial prefrontal cortex and amygdala of aged Wistar rats (Sotoudeh et al. 2020). Dendritic spine classification revealed an increased presence of stubby spines in infralimbic and prelimbic areas of the prefrontal cortex. In a study conducted in aged rats subjected to behavioral stress, thin and stubby spines from the prefrontal cortex of aged rats were more susceptible to stress, while mushroom spines remained even after the stimuli, probably due to reduced dendritic spine plasticity (Bloss et al. 2011). The presence of stubby spines induced by the KD could represent a plausible explanation for the enhanced synaptic plasticity and performance in the Y-maze test; however, further experiments are needed to understand in detail the effects of the KD on dendritic spine dynamics of aged mice. It is important to note that the preserved basal transmission observed in electrophysiological assays is not contradictory with the higher dendritic arborization and spine density in KD-fed animals, as brain circuits are endowed with several homeostatic mechanisms allowing to maintain synaptic strength in a functional range. It is the increase in LTP which reveals an improvement in the plastic properties of synapses that might underlie the enhanced learning abilities of KD-fed mice.

The hallmarks of aging encompass a range of biological processes that lead to functional decline in an organism, and include the loss of proteostasis (López-Otín et al. 2023; Hipp, Kasturi, and Hartl 2019; Kaushik and Cuervo 2015). Indeed, neurodegenerative diseases display an accumulation of aberrant proteins as a common feature. Interestingly, therapies that promote the activation of mechanisms involved in protein folding enhance cognitive function in aging mice (Bobkova et al. 2015). Recently, Hafycz and cols, administrated a chemical chaperone 4-phenyl butyrate into aged mice and observed improved learning through increased p-CREB, a key protein involved in memory consolidation (Hafycz, Strus, and Naidoo 2022). In this work, we evaluated the changes in the cortical proteome of aged mice after a long-term administration of a cyclic KD, as optimized in (Newman et al. 2017). Proteomics data showed that a cyclic KD modifies predominantly the presynaptic proteome and regulates pivotal biological processes involved in synaptic function, such as actin cytoskeleton dynamics, the cAMP-mediated signaling pathway and synaptic vesicle cycling. Consistently, in 2010, a proteomics study performed in rat synaptosomes found that aging downregulated the expression of key proteins involved in synaptic vesicle trafficking (Van Guilder et al. 2010), suggesting that a KD might be involved in synaptic vesicle regulation. Compelling evidence points out that the cAMP-mediated signaling pathway plays a key role in LTP and memory consolidation (Bernabeu et al. 1997; Murphy and Segal 1997; Bach et al. 1999). There is no clear consensus about the levels of cortical cyclic AMP during the aging process. Whereas some reports indicate a pathway disinhibition in the brain cortex that compromises working memory (Vandesquille et al. 2013; Ramos et al. 2003), others suggest a reduction in cAMP in the prefrontal cortex of rodents (Zimmerman and Berg 1974; Puri 1981; Hara et al. 1992; Wruck and Adjaye 2020). Our findings are in line with the latter studies, as shown by an increase in adenylate cyclase 5 (Adcy5), Protein Kinase A catalytic subunit (Prkacb), Phosphodiesterase 10a (Pde10a) and Phosphodiesterase 1b (Pde1b) for the KD group. The synaptic vesicle cycle consistently emerges as an annotation pathway observed in the GO characterization. Interestingly, actin-related proteins form part of that group in the GO terms, such as Complexin 1 (Cplx1), Annexin V (Anxa5) and Rac Family Small GTPase 1 (Rac1). PKA kinase activity plays a critical role in synaptic plasticity (Havekes et al. 2012; Park et al. 2014) mediating synaptic vesicle trafficking and release through synapsin 1 phosphorylation (Patzke et al. 2019), whereas cAMP/PKA signaling has been associated with the regulation of actin filaments (Nadella et al. 2009; Shabb 2001). Thus, further experiments focused on PKA substrate phosphorylation could reveal new molecular mechanisms associated with enhanced synaptic activity efficiency induced by a KD.

Proteomics data information permitted an in-depth investigation into the molecular mechanisms by which the diet may regulate brain function. Thus, we prepared whole protein extracts from brains of aged mice which had been fed with a cyclic KD for 4 months. Microtubules are an important structural support for synaptic plasticity, and microtubule associated proteins (MAPs) regulate dendritic differentiation and maintenance (Conde and Cáceres 2009). Intriguingly, western blot analysis revealed an increase in MAP2 phosphorylation for the KD group. The beneficial effects of a KD on brains were first applied clinically to treat refractory epilepsy which is characterized by an aberrantly augmented excitatory tone. At the molecular level, Sánchez and cols, demonstrated in an *in vitro* culture of hippocampal neurons, that MAP2 phosphorylation decreases in seizure-like activity induced by kynurenic acid, which is related to increased microtubule rigidity and hyper-stabilization (Sánchez et al. 2001). In contrast, we observed increased MAP2 phosphorylation. This modification promotes more dynamic MAP2 association with microtubules, contributing to the versatile organization of tubulins in dendrites of aged mice. This could partially explain the reduced occurrence of seizure episodes and the increased dendritic arborization observed in pyramidal neurons of the KD group. When PKA protein levels were compared between groups, no significant differences were registered, although an increasing trend was observed in the KD condition. Thus, we postulate that changes in the cAMP-pathway induced by the KD modify the activity, rather than the amount, of PKA. Supporting this idea, we also observed up-regulation of the phosphodiesterases Pde1b and Pde10a in the KD condition, likely suggesting that the fine-tuning that controls the activity of cAMP levels and PKA activity is much more complex than by simply increasing the relative abundance of the catalytic subunit of PKA. Thus, more studies are needed to unravel this novel finding associated with the KD. Nevertheless, although PKA expression was not substantially upregulated by the KD, we confirmed the activation of the signaling pathway, as brain derived neurotrophic factor (BDNF), a canonical target of this route, was overexpressed in the KD group. BDNF activation is mediated by p-CREB; thus BDNF regulates synaptic plasticity and structural changes in dendritic spines, promoting learning and memory processes (Gómez-Palacio et al. 2013; Tanaka et al. 2008). Additionally, it is well known that BDNF and neurotrophic factor signaling is impaired in the aging brain (Mattson, Maudsley, and Martin 2004; Tapia-Arancibia et al. 2008; Mattson and Arumugam 2018). Notably, here we report that the KD induces BDNF expression in the brain cortex of aged mice, even when it is administered later in life, which is consistent with the improvements in spatial working memory and cognition that we observed. Neuronal BDNF regulation in response to a KD was reported earlier when Hu and cols found that BHB, the main ketone body produced in the KD, induced BDNF expression in the hippocampus and primary cultures of hippocampal neurons treated with this substance. Interestingly, they also observed an increase in the cAMP/PKA signaling pathway, the main regulator of BDNF expression (Hu et al. 2018).

Nevertheless, this is the first report showing that systemic production of ketone bodies induces cortical BDNF expression in aged brains. Our biochemical approach also confirmed that PKA-dependent signaling was indeed enhanced after application of the KD. We determined *in vivo* and *in vitro* that this nutritional intervention activates PKA, using an antibody against phosphor-epitopes that recognizes the consensus site RRXS*/T as a surrogate marker. Primary culture experiments showed that we can reproduce *in vitro*, findings observed *in vivo*.

Moreover, we determined that a 130 kDa protein was significantly phosphorylated by PKA *in vitro* and *in vivo*. Although the identity of the proteins that are more robustly phosphorylated remains unknown, these results merit further studies to address the identity and sites of phosphorylation of PKA substrates triggered by the KD. Importantly, we detected CREB phosphorylation in cultures treated with BHB, which confirms and reinforces the bioinformatics findings acquired from mass spectrometry analysis. Finally, as a proof of concept, we employed AKAR4, a widely used PKA activity-based biosensor in COS7 cells (Depry et al. 2011). When BHB was added, a strong FRET ratio increase was observed, suggesting the rapid activation of PKA kinase in response to this metabolite.

The present study presents some limitations which are related to the fact that we studied cortical areas in our experimental design, setting aside changes that the KD could elicit in the hippocampus, beyond the LTP effects reported here. We also did not delve into the phosphoproteins regulated by PKA; however, we certainly consider that exploring this downstream signaling pathway in more detail merits further analysis.

In summary, our data provides new insights into molecular mechanisms and biological processes that a cyclic KD regulates in brain function, an aspect that has been understudied in the field of aging. In addition, we revealed here that a KD has the potential to modify brain function and motor activity in aged mice, even when administrated later in life. This study also proposes new mechanisms by which the administration of a cyclic KD improves memory and neuronal function in aging that had not been discovered previously. Specifically, a KD induces changes in the proteome landscape of cortical synapses that directly impact the structure and function of synaptic organization, proposing a scenario whereby ketone bodies (specifically BHB) play a crucial role not only as an energy metabolite, but also as a signaling metabolite (Newman and Verdin 2017).

## Supporting information

Candidates_DA3

## Acknowledgements

We acknowledge the support of instrumentation for the TripleTOF 6600 from the NIH shared instrumentation grant 1S10 OD016281 (Buck Institute). Thiswork was supported by the following grants: ANID/Fondecyt/1220414 and ANID/FONDAP/15150012 to CG-B; ANID Scholarship 75200183 to DA, IMPACT (FB210024, EA, UW), Fondecyt Grant (3190897 DC), ACDicyt (5392202MMÑ,BM), NIH Grant (AG06733, BS) and NIH Grant (1S10 OD016281 01, JCN).We thank Professor Michael Handford for English editing

## Supplemental information

**Figure S1:**
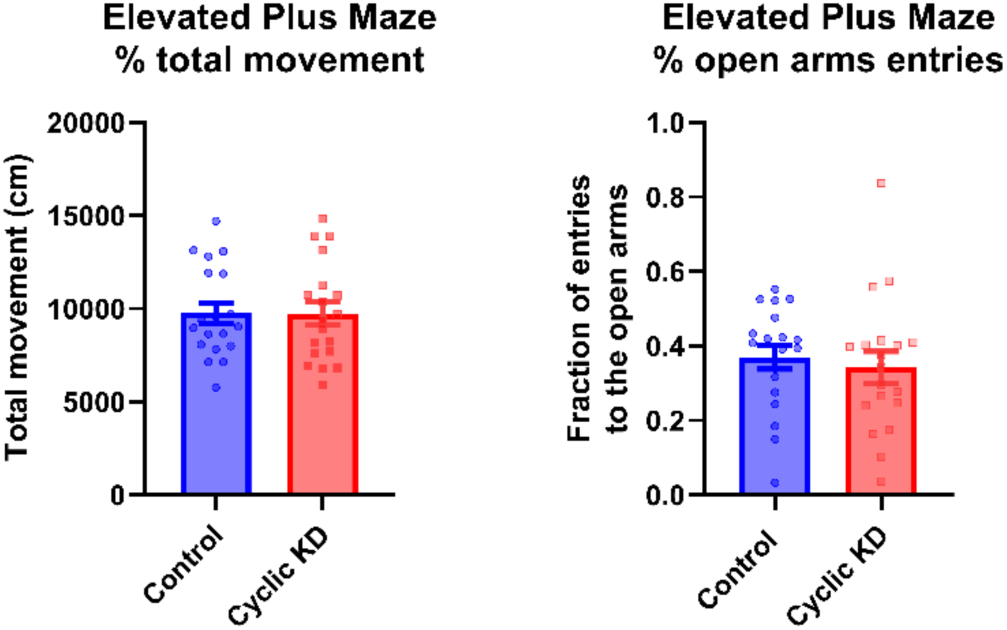
Complement to figure 2. Elevated plus maze test of aged mice fed with a control or a cyclic KD. (A) Mice were subjected to an elevated plus maze to assess anxiety-like behavior and total movement during the entire test. (B) The number of entries to the open arms was calculated and expressed as the fraction of total entries (n=19 per group). p>0.05, t-student test. Error bars show the SEM.

**Figure S2:**
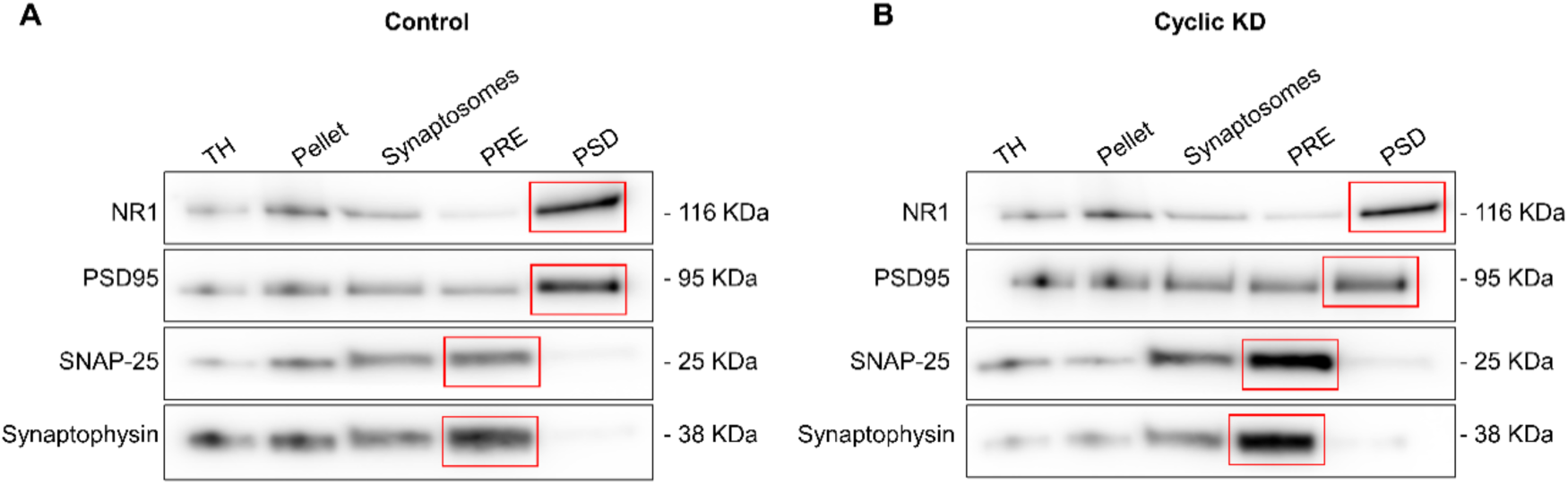
Complement to figure 5. Synaptosomes preparation from prefrontal cortex of aged mice fed with control and cyclic KD. **(A, B).** Prefrontal cortices from aged mice fed with control and cyclic KD by 52 weeks were collected in control week and processed for synaptosomes preparation. Total homogenates (TH) and pellet were collected for a complete characterization. Synaptosomes were separated into enriched presynaptic and postsynaptic fractions (here referred as PRE and PSD respectively) for subsequent western blot characterization. NR1, PSD95, SNAP-25 and Synaptophysin antibodies were used for immunodetection of postsynaptic and presynaptic proteins, showing enrichment in the respective fractions (highlighted in red boxes).

**Figure S3:**
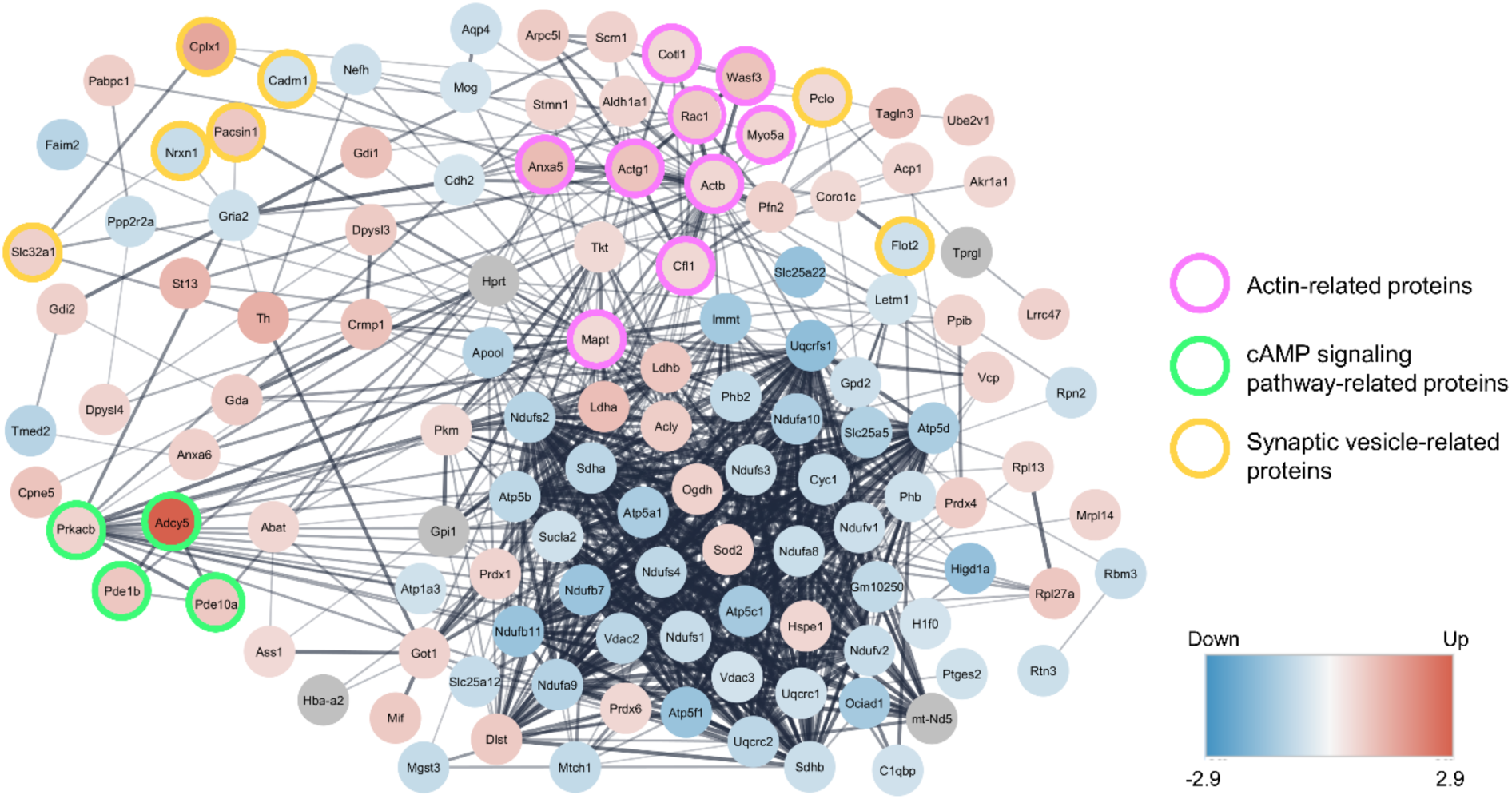
Complement to figure 5. Differentially expressed proteins found in cortical synaptosomes of aged mice fed with a long-term cyclic KD. Total differentially expressed proteins were analyzed using the STRING database. A cluster of cAMP-mediated signaling pathways showed substantial upregulation in response to the KD, including adenylate cyclase 5 (Adcy5), cAMP-dependent protein kinase catalytic subunit beta (Prkacb), phosphodiesterase 1b (Pde1b) and phosphodiesterase 10a (Pde10a) (light green circles). Actin-related proteins and synaptic vesicle-related proteins are highlighted in magenta and light orange, respectively. The confidence score for interactions was 0.4.

**Figure S4:**
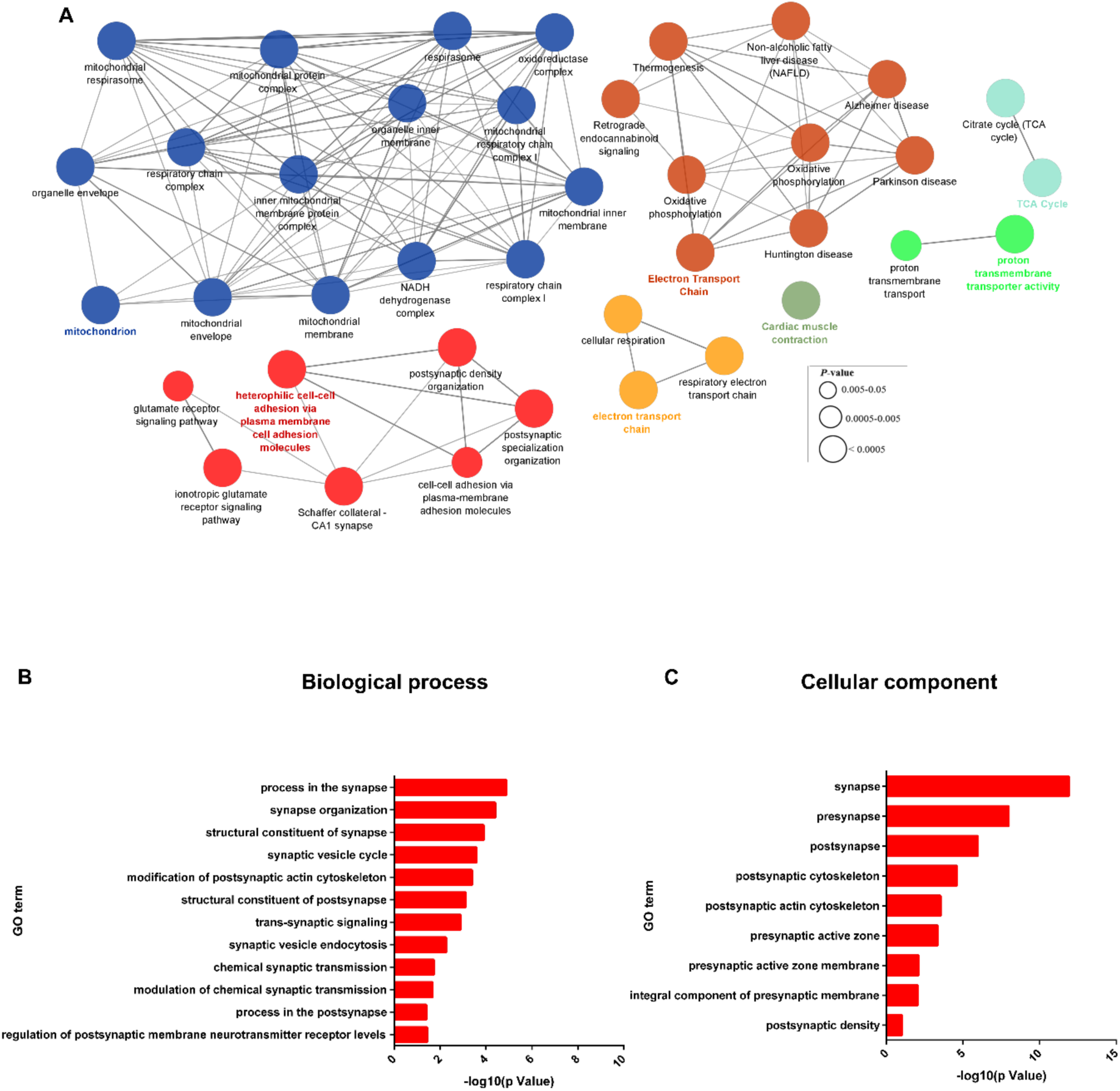
Complement to figure 5. Bioinformatics analysis revealed mitochondrial function and the synaptic vesicle cycle as biological processes regulated in aged mice fed with a long-term cyclic KD. (A) The cortical synaptosome proteome of mice fed with a cyclic KD for 52 weeks showed mitochondria as a biological process that was downregulated as a result of the treatment. Samples were collected in a control week (high presence of carbohydrates), demonstrating the rapid response to the metabolic switch induced by diet cycling. (B-C) Characterization of the cortical synaptic proteome using SynGO revealed the synaptic vesicle cycle as a representative biological process regulated by the long-term administration of a cyclic KD (red arrows).

## Methods

### Figure panels

Figure plots were designed in GraphPad Prism (Version 8.2.1(279) and 9.4.1(458)).

### Mouse strain, housing, and diet administration

Male C57BL/6 mice were acquired from Jackson Laboratory at 18 months of age and maintained in a facility with a 12 h light/dark cycle at 25°C with food and water provided *ad libitum*. All animal procedures were approved by the Bioethics Committee of the Faculty of Medicine of Universidad Mayor, Chile. Mice were housed in groups, with up to 5 mice per cage. Cages contained absorbent paper chip bedding which was changed every week. Enrichment was provided in the form of a nestlet. Experimental diets were changed weekly. Control food was obtained from Envigo (TD.150435) containing: 10% protein, 13% fat, 77% carbohydrates, whereas the cyclic ketogenic diet (KD; TD.160153) contained 10% protein and 90% fat. Fat sources are Crisco, cocoa butter, and corn oil. The detailed food composition of the two experimental diets is described in (Newman et al. 2017).

### Metabolic parameter measurements

Blood was collected via distal tail-snip (10 - 20 µL) for glycemia and β- hydroxybutyrate measurements using Abbott FreeStyle Optium strips. Blood draws were taken early in the morning once a week during 12 consecutive weeks. Body weight and food consumption were registered every week throughout the experiment.

### Elevated plus maze

Th elevated plus maze (Rodgers and Dalvi 1997) is a task to measure anxiety-like behavior, and consists of two platforms (78 cm high x 5 cm wide) arranged in a cross-shaped configuration. One platform is open whereas the other has a continuous wall, 10 cm high. Mice were allowed to explore the platforms for 10 minutes. Covered distance and permanence time in open or closed platforms were recorded and measured using ANY-maze software (version 7.01, Stoelting Co., USA).

### Open field

The open field test was performed using a 40 cm cubic box. Mice were individually placed at the center of the platform. Spontaneous activity was recorded for 15 minutes as described in (Chew et al. 2015) and then analyzed with ANY-maze software. Between trials, the arena was cleaned with 30% ethanol to remove odors. Total distance traveled and total distance spent at the center were calculated. An imaginary 20 x 20 cm center zone was used to define the center:total distance ratio.

### Novel object recognition

The novel object recognition test was performed using an uncovered 40 cm cubic box, and consisted of familiarization and probe steps on two consecutive days (Antunes and Biala 2012). On the first day (familiarization), the animals were placed in the arena and left to explore for 10 min. Subsequently, 2 identical objects were placed near (5 cm) to corners, the mice were placed back inside the box looking at the wall opposite to the objects, and then tracked for 10 min. Next day (probe), animals were placed in the arena to explore for 5 min then removed. Subsequently, one object from the day before (known object) and a new object, different in shape, height and color, were placed near (5 cm) to corners. Mice were then put back in the box looking at the wall opposite to the objects, and tracked for 10 min.

All objects had a 5-cm diameter cylindrical shape base, and the exploration of the objects was determined to within 2.5 cm of the objects. All tracking was performed by ANY-maze software. Between trials, the arena was cleaned with 30% ethanol to remove odors.

### Y-maze

The Y-Maze test was performed using a maze with 3 arms (8 cm wide, 35 cm long and walls of 20 cm height) connected at 120° angles, forming a Y. Each arm was assigned A, B or C (Kraeuter and Guest 2019). Each animal was placed in the arm (A) looking towards the central node of the maze and tracked for 8 min by ANY-maze software. Between trials, the arena was cleaned with 30% ethanol to remove odors. Alternation was defined as visits to 3 different arms (ABC, ACB, BCA, BAC, CAB, CBA) and % spontaneous alternation was determined by the formula: % spontaneous alternation = (Total alternations / (Total arms entries - 2)) x 100.

### Barnes Maze

In the Barnes maze, mice were placed on a circular arena (92 cm diameter and 72.5 cm above the floor) containing 20 equally spaced holes on the edge (5 cm diameter and 7.5 cm apart from each other) (Sunyer et al. 2007). Visual cues were placed around the arena for spatial navigation. Before each trial, mice were placed for 10 seconds inside a cylindrical container in the center of arena. Upon removal of the cylinder, the animals were exposed to an aversive sound (beep of a timer). The time the animal took to find and enter the target hole leading to an escape box was recorded as primary and total latency, respectively. The protocol consisted of 3 phases: adaptation, acquisition, and test. During adaptation, the mice were gently guided into the escape box by the experimenter and allowed to stay there for 2 minutes. The acquisition phase consisted of 4 trials per day, during 4 consecutive days (day 1 - 4). During the acquisition phase, each animal was allowed to explore the maze for 3 minutes or until entering the escape box. After the mouse had entered the box, the sound was immediately turned off and the animal allowed to stay there for 1 minute. If the mouse did not enter the escape box within 3 minutes, the experimenter guided the animal into the box, allowing it to stay there for 1 minute. Each mouse was then placed in its cage until the next trial (with 15-minute intervals between trials). Between trials, the arena and wall were cleaned with 30% ethanol to remove odors. Primary and total latency were measured. The test was conducted at days 5 and 12 to assess short- and long-term spatial memory, respectively. The target hole without the escape box was maintained in the same position used in the acquisition period. The test was performed for 90 seconds, and the latency recorded. The primary latency from days 1 to 4 was measured as learning curve, at day 5 as short-term memory, while at day 12 as long-term memory.

### Rotarod

The test was performed using Rotarod LE8200 Panlab Harvard Apparatus (Barcelona, Spain) on two consecutive days (Deacon 2013). On the first day (training), all animals were placed on rotating lines with a constant speed of 4 rpm for 1 minute, three times. Later, on the second day (test), all mice were placed on rotating lines, three times, with acceleration set from 4 to 40 rpm over 5 minutes, and the time noted when the mice fell off.

### Hippocampal slices for electrophysiology

Transversal slices were prepared from 25-27 m.o mice fed with a control or a cyclic KD. Animals were deeply anesthetized with isoflurane and transcardially perfused with ice-cold oxygenated (95% O_2_, 5% CO_2_) N-methyl-D-glucamine (NMDG) based solution for a better cell preservation in slices (Ting JT, Daigle TL, Chen Q). After perfusion, the brain was quickly removed and submerged in ice-cold oxygenated (95% O_2_, 5% CO_2_) NMDG solution containing (in mM): 93 NMDG, 2.5 KCl, 1.2 NaH_2_PO_4_, 30 NaHCO_3_, 20 HEPES, 25 glucose, 2 thiourea, 5 sodium ascorbate, 3 Na-pyruvate, 0.5 CaCl_2_, and 10 MgSO_4_, pH 7.4 adjusted with HCl. Slices were cut in this solution at a thickness of 300 µm using a Leica vibratome. After slicing, the tissue was incubated for 10 min at 32°C in NMDG buffer solution. Then, slices were transferred into a storage chamber kept at RT in artificial cerebrospinal fluid (ACSF) containing (in mM): 124 NaCl, 2.5 KCl, 1.2 NaH_2_PO_4_, 24 NaHCO_3_, 5 HEPES, 12.5 glucose, 2 CaCl_2_ and 2 MgSO_4_, pH 7.4 (95% O_2_, 5% CO_2_).

### Electrophysiology

Extracellular recordings in hippocampal slices were performed as described in (Contreras et al. 2022). Field excitatory postsynaptic potentials (fEPSPs) were evoked by applying electrical stimulation delivered by an A360 stimulus isolator (WPInc, Sarasota, FL, USA), using bipolar concentric electrodes (200 μm diameter; FHC Inc., Bowdoinham, ME, USA) on Schaeffer collateral-commissural fibers and recorded with glass microelectrodes (1–2 MΩ) filled with ACSF from the stratum radiatum of the hippocampal CA1 region. Test pulses (0.2 ms) were applied every 15 s, and the current was adjusted to evoke 50% to 60% of the maximal response. After recording a stable baseline for at least 20 min, LTP was induced by a theta burst stimulation (TBS, 5 trains of 10 bursts at 5 Hz each; 1 burst = 4 pulses at 100 Hz). In all experiments, the fEPSP recordings were maintained for 60 min after initiating the TBS. The synaptic responses were quantified as the initial slope of fEPSPs and plotted as a percentage of change, referring to the initial slope measured during the baseline recording before TBS.

For input-output and paired-pulse experiments we used glass microelectrodes filled with ACSF for both stimulation and recording. Input-output curves were obtained by applying stimuli of increasing intensities (from 0.1 to 1.0 mA, with 0.1 mA steps; 2 recordings per stimulus amplitude), after reaching a stable transmission baseline of 5–10 min. Paired pulse facilitation was assessed by delivering 2 stimulation pulses separated by 25, 50, 100, 150, 200 and 500 ms. The current intensity was set to evoke a first response of 50-60% of the maximum obtained in the input-output protocol. Paired-pulse facilitation was calculated by dividing the slope of the second response to that of the first. Data was analyzed using IgorPro 6.3 (Wavemetrics) and statistical tests were performed with GraphPad prism software (Dotmatics).

### Golgi staining

Whole brains were collected, washed in PBS 1X, and stained using the Hito Golgi-Cox OptimStain Kit (Hitobiotec, Corp, Kingsport, TN) following the manufacturer’s instructions. Briefly, brains were immersed in the impregnation solution (by mixing solution 1 and 2) and stored at room temperature avoiding light exposure for 14 days. Then brains were transferred into solution 3, which was replaced 12 hours later and kept at 4°C in the dark for 72 hours. Tissues were quickly frozen with isopentane previously chilled at -80°C for 1 minute, and then embedded in a 30% sucrose solution at low temperature in a cryostat (Microm HM525, Thermo Scientific, MA, USA). Using the Allen mouse brain atlas as reference, coronal sections of 300 µm containing the prefrontal cortex were sliced, mounted onto gelatin-coated slides, and stained in a mixture of solution 4, solution 5 and distilled water for 10 minutes. Finally, samples were covered with a coverslip using xylene-based resinous mounting medium.

### Morphometric analysis

Coronal sectioning at 150 μm was performed to analyze dendritic morphology. The morphometric analysis of pyramidal neurons was restricted to the following stereotaxic coordinates (in mm): for the medial Prefrontal Cortex (mPFC) cortex from 5.78 to 5.14 interaural, and from 1.98 to 1.34 Bregma (Paxinos and Watson, 1998). Golgi-stained images were obtained with an Olympus CX41 epifluorescence microscope with a Plan Fluor objective/numeric aperture: 20x/1.25, 40x/0.65 and 100x/1.25. Pyramidal neurons were defined by the presence of basilar dendrites, with a distinctive, single apical dendrite and dendritic spines (Spruston, 2008). The observer, blind to experimental conditions, selected randomly at least 10 neurons from each animal that fulfilled the following selection criteria: 1) absence of truncated dendrites, 2) consistent and dark impregnation across the entire dendritic field, and 3) spatial separation from neighboring impregnated neurons to avoid overlap. Camera lucida tracings from selected neurons (500x, BH-2, Olympus Co., Tokyo, Japan) were performed and then scanned (eight-bit grayscale TIFF images with 1,200 dpi resolution; EPSON ES-1000C) along with a calibrated scale for subsequent computerized image analysis. For morphometric analysis of digitized images, Image J software (National Institutes of Health) was used.

For the dendritic spine analysis, spines were visualized with a 100x/1.25 Plan Fluor objective. Control (14) and 20 pyramidal neurons, and 1508 control and 1744 keto spines were measured, considering primary dendrites as those starting from the origin of the soma, and secondary dendrites as those that originate from a branch point in the primary dendrites. The spines were counted in 5 µm segments along a 50 µm stretch of the dendrite. Dendritic spines were classified under the microscope at different focal planes, by a single rater, blinded to the experimental groups, into three shape categories (Harris et al., 1992): filopodia/thin, stubby, and mushroom/branched spines. Thin spines had a greater length than neck diameter and similar head and neck diameters. Stubby spines had neck diameters that were similar to the total spine length. For mushroom-shaped spines, the diameter of the head was much greater than the diameter of the neck. Branched spines had more than one head emerging from a single neck originating from the dendrite.

### Synaptosomes preparation

Prefrontal cortex tissue was collected from 8 C57BL/6 mice, 4 of which were on a control diet for 52 weeks, and 4 of which were on a cyclic KD diet for 52 weeks. For each tissue sample, a presynaptic fraction and a postsynaptic fraction were obtained, giving a total of 16 samples. Synaptosomes were obtained according to (Dosemeci et al. 2006). Briefly, samples were homogenized with a buffer containing 0.32 M sucrose, 5 mM Tris-HCl pH 7.4, 1 mM PMSF, and a cocktail of protease and phosphatase inhibitors, using Tissuelyser II (Qiagen) and steel beads of 5 mm diameter. The homogenate was centrifuged at 1000 g for 2 minutes and the supernatant was collected by centrifugation at 12000 g for 20 minutes. Pellets were resuspended in a buffer with 0.32 M sucrose, 0.5 mM EGTA and 5 mM Tris-HCl pH 7.4 and then placed on the top of a gradient with 1.2 M/1 M sucrose. Samples were centrifuged at 250000 g for 60 minutes in a swinging bucket rotor of the ultracentrifuge (Beckman Coulter Optima L-60). Then, total synaptosomes from the 1.2 M/1 M sucrose interphase were collected, diluted 1:5 with buffer containing 0.5 mM EGTA, 10 mM DTT, 5 mM Tris-HCl pH 8.1 and a cocktail of protease and phosphatase inhibitors and then incubated for 30 minutes at 4°C. Samples were centrifuged at 33000 g for 30 minutes and a supernatant enriched in presynaptic proteins was called “PRE”, while the pellet enriched in post synaptic densities was called “PSD”. The presynaptic fractions were precipitated overnight with pre-chilled acetone and washed with cold acetone the next day. Pellets were air dried and re-suspended in 5 mM Tris-HCl buffer, pH 7.4 with 0.5% SDS. The postsynaptic fractions were centrifuged in a buffer containing 0.32 M sucrose, 0.5 mM CaCl_2_, X-100 Triton, 2 mM DTT and 10 mM Tris, pH 8.1. These pellets were also re-suspended in 5 mM Tris, pH 7.4 with 0.5% SDS.

### Protein digestion

A Bicinchoninic Acid protein assay was performed for each of the synaptosome samples, and a 30 µg aliquot was used for tryptic digestion for each of the 16 samples. Protein samples were added to a lysis buffer containing a final concentration of 5% SDS and 50 mM triethylammonium bicarbonate (TEAB), pH ∼7.55. The samples were reduced in 20 mM dithiothreitol for 10 minutes at 50°C, subsequently cooled at RT for 10 minutes, and then alkylated with 40 mM iodoacetamide for 30 minutes at RT in the dark. Samples were acidified with a final concentration of 1.2% phosphoric acid, resulting in a visible protein colloid. Methanol (90%) in 100 mM TEAB was added at a volume of 7 times the acidified lysate volume. Samples were vortexed until the protein colloid was thoroughly dissolved in the 90% methanol. The entire volume of the samples was spun through the micro-S-Trap columns (Protifi) in a flow-through Eppendorf tube. Samples were spun through in 200 µL aliquots for 20 seconds at 4000 g. Subsequently, the S-Trap columns were washed with 200 µL of 90% methanol in 100 mM TEAB (pH ∼7.1) twice for 20 seconds, each at 4000 g. S-Trap columns were placed in a clean elution tube and incubated for 1 hour at 47°C with 125 µL of trypsin digestion buffer (50 mM TEAB, pH ∼8) at a 1:25 ratio (protease:protein, wt:wt). The same mixture of trypsin digestion buffer was added again for an overnight incubation at 37°C.

Peptides were eluted from the S-Trap column the following morning in the same elution tube as follows: 80 µL of 50 mM TEAB was spun through for 1 minute at 1000 g, then 80 µL of 0.5% formic acid was spun through for 1 minute at 1000 g. Finally, 80 µL of 50% acetonitrile in 0.5% formic acid was spun through the S-Trap column for 1 minute at 4000 g. These pooled elution solutions were dried in a speed vac and then re-suspended in 0.2% formic acid.

### Desalting

The re-suspended peptide samples were desalted with stage tips containing a C18 disk, concentrated and re-suspended in aqueous 0.2% formic acid containing “Hyper Reaction Monitoring” indexed retention time peptide standards (iRT, Biognosys).

For mass spectrometry, samples were analyzed by reverse-phase HPLC-ESI-MS/MS using an Eksigent Ultra Plus nano-LC 2D HPLC system (Dublin, CA) with a cHiPLC system (Eksigent) which was connected directly to a quadrupole time-of-flight (QqTOF) TripleTOF 6600 mass spectrometer (SCIEX, Concord, CAN). After injection, peptide mixtures were loaded onto a C18 pre-column chip (200 µm x 0.4 mm ChromXP C18-CL chip, 3 µm, 120 Å, SCIEX) and washed at 2 µl/min for 10 min with the loading solvent (H_2_O/0.1% formic acid) for desalting. Subsequently, peptides were transferred to the 75 µm x 15 cm ChromXP C18-CL chip (3 µm, 120 Å; SCIEX), and eluted at a flow rate of 300 nL/min with a 3 h gradient using aqueous and acetonitrile solvent buffers.

### Data-independent acquisitions

For quantification, all peptide samples were analyzed by data-independent acquisition (DIA, e.g. SWATH), using 64 variable-width isolation windows (see Supplementary table 1) (Collins et al. 2017). The variable window width was adjusted according to the complexity of the typical MS1 ion current observed within a certain m/z range using a DIA ‘variable window method’ algorithm (narrower windows were chosen in ‘busy’ m/z ranges, whilst wider windows were chosen in m/z ranges with few eluting precursor ions). DIA acquisitions produce complex MS/MS spectra, which are a composite of all the analytes within each selected Q1 m/z window. The DIA cycle time of 3.2 sec included a 250 msec precursor ion scan followed by 45 msec accumulation time for each of the 64 variable SWATH segments.

### Mass-spectrometric data processing, quantification, and bioinformatics

The DIA/SWATH data was processed in Spectronaut version 14.2.200619.47784 (Biognosys) using a custom in-house spectral library. Data extraction parameters were selected as dynamic, and non-linear iRT calibration with precision iRT was selected. Identification was performed using a 1% precursor and protein q-value, and iRT profiling was selected. For the DIA/SWATH MS2 data sets, quantification was based on the XICs of 3-6 MS/MS fragment ions, typically y- and b-ions, matching to specific peptides present in the spectral libraries. Peptide abundances were obtained by summing precursor abundances and protein abundances by summing peptide abundances. Interference correction was selected and local normalization was applied. Differential protein abundance analysis was performed using paired t-test, and p-values were corrected for multiple testing, specifically applying group-wise testing corrections using the Storey method (Burger 2018; Storey 2002). Significantly changed proteins were accepted with ≥ 2 unique peptides, q-value < 0.05 and absolute log_2_(fold change) ≥ 0.58.

### Data availability

The mass spectrometric raw data and complete datasets have been uploaded to the MassIVE repository of the Center for Computational Mass Spectrometry at UCSD and can be downloaded using the following link: https://massive.ucsd.edu/ProteoSAFe/dataset.jsp?task=b07ee0b4d42c46029d096 9d147a63a0c (MassIVE ID number: MSV000092626; ProteomeXchange ID: PXD044458).

[Note to the reviewers: To access the data repository MassIVE (UCSD) for MS data, please use: Username: MSV000092626_reviewer; Password: winter].

### Pathway and network analysis

Proteomics data was analyzed with R studio software (version 1.3.1093) using the clusterProfiler package (Guangchuang et al. 2012). Pathway enrichment characterization was performed with ClueGO package, version 2.5.3, in Cytoscape (https://cytoscape.org/), version 3.7.1 (Shannon et al. 2003; Bindea et al. 2009) and was referenced from the following databases: GO Biological Function, GO Cellular Compartment, Kegg Pathways, WikiPathways, and Reactome Pathways including pathways with experimental evidence (EXP, IDA, IPI, IMP, IGI, IEP). For statistical cutoff, the Benjamini-Hochberg test was used with adjusted p-values<0.01 by right-sided hypergeometric testing. Edges connecting pathways were drawn based on kappa scores >40%. Kappa scores serve as a measure of inter-pathway agreement between observed proteins, indicating whether the agreement between pathways is higher than what would be expected by chance, considering shared proteins. Pathways that share the same color indicate ≥50% of similarity in GO terms. Functional annotation of synaptic proteins was performed with the SynGO database (https://www.syngoportal.org; Koopmans et al. 2019). Additionally, the STRING package in Cytoscape was used to study protein-protein associations (von Mering et al. 2005).

### Western Blot

Whole protein extracts from prefrontal cortices were processed for western blot characterization. Proteins were quantified using the Qubit 4 fluorometer (ThermoFisher Scientific, MA, USA) and resolved by 10% SDS polyacrylamide gels electrophoresis. Samples were transferred onto PVDF membranes and blocked with blocking buffer containing 5% bovine serum albumin, 1X TBS-T (1X Tris-Buffered Saline, 0.1% Tween) for 1 hour at RT. Then, membranes were probed over night at 4°C with agitation using the following antibodies: Anti p-MAP2 305, kindly donated by Dr. Jesús Ávila (CBMSO, Madrid, Spain) (Sánchez et al. 2001), anti MAP-2 (#AB5622-I Sigma Aldrich), anti Gria2 (#SAB4300535 Sigma), anti PKA β-subunit (#ab187515 Abcam), anti BDNF (N20) (#SC546 Santa Cruz), anti p-PKA substrates (RRXS*/T*) (#9624 Cell signaling) and anti GAPDH (#MAB374 Merck). Membranes were washed 3 times with 1X TBS-T buffer and incubated with anti-mouse (#AB_2340770) or anti rabbit (#AB_10015282) HRP secondary antibodies (Jackson ImmunoResearch Laboratories, PA, USA) for 1 hour at RT. Proteins were detected using the SuperSignal West Pico PLUS Chemiluminescent Substrate kit (#34580, ThermoFisher Scientific, MA, USA) and images were taken with a Uvitec Cambridge (Uvitec Cambridge, Cambridge, UK) chemiluminescence imaging system, using Alliance software version 16.06. Finally, bands were quantified using ImageJ software.

### Primary culture, cell lines and *in vitro* β-hydroxybutyrate treatments

E18.5 cortical neurons were obtained from cortices of C57BL/6 embryos as described in (Wilson, et al. 2020). Briefly, cortices were dissected and treated with 0.25% trypsin and 1X DNAse for 25 minutes at 37°C. Tissue was gently disaggregated and neurons plated for 90 minutes with DMEM 10% HS medium in P6 well plates previously treated with 1 mg/ml poly-L-lysine (Sigma Aldrich) at a density of 300,000 cells/cm^2^. Then, DMEM 10% HS was replaced by neuronal maintenance medium with N2 supplement (Gibco), and neurons were cultured until 28 DIV at 37°C with 5% CO_2_. At day 27, DIV neurons were treated with 5 mM R-β-hydroxybutyrate (#54920, Sigma Aldrich) in low (2.5 mM) glucose media for 24 hours to reproduce the physiological effects of the metabolic switch elicited by the KD. At day 28, DIV neurons were treated with BHB for 1 hour and processed for whole protein extraction. COS7 cells were cultured in DMEM 10% FBS and transfected with the AKAR4 plasmid (#61619, addgene) for 24 hours. Then, media concentrations were reduced to 2.5 mM glucose and 5 mM BHB and incubated for 1 hour. After treatments, cells were fixed with 4% paraformaldehyde for 20 minutes, mounted on slices, and visualized with a confocal microscopy Zeiss LSM 710 using the 40x oil immersion objective. Images were processed with ImageJ.

## Notes

### Competing Interest Statement

The authors have declared no competing interest.

